# Cell-type specific responses to single-pulse electrical stimulation of the human brain

**DOI:** 10.1101/2024.11.24.625077

**Authors:** Rhiannon L. Cowan, Tyler S. Davis, Edward M. Merricks, Bornali Kundu, Ben Shofty, Shervin Rahimpour, Catherine A. Schevon, John D. Rolston, Elliot H. Smith

**Affiliations:** Department of Neurosurgery, University of Utah, Salt Lake City, UT 84132, USA; School of Medicine, University of Missouri, Columbia, MO 65212, USA; Department of Neurology, Columbia University, NY 10032, USA; Department of Neurosurgery, Mass General Brigham, Harvard Medical School, Boston, MA 02115, USA

**Keywords:** neuromodulation, epilepsy, Corticocortical evoked potential (CCEP), single-unit activity, Single-pulse electrical stimulation (SPES), Responsive Neurostimulation (RNS)

## Abstract

Neuromodulation techniques, such as deep brain stimulation, intraoperative brain mapping, and responsive neurostimulation, use electricity to alter brain activity. Despite daily clinical use in thousands of patients, it remains fundamentally unknown how human neurons respond to intracranial stimulation.

We address this question at a basic level by characterizing neuronal cell-type specific firing rate responses to single pulses of electrical stimulation of the human brain. We carried out broadly distributed stimulation in 30 patients undergoing neuromonitoring for epilepsy while recording from isolated neurons on microwires implanted into the medial temporal and frontal lobes.

Out of a total of 228 recorded units, 16.2% (N = 191) were classified as interneurons and 83.8% (N = 37) were classified as principal cells, using a threshold clustering method, based on intrinsic waveshape characteristics. To see how stimulation affected neuronal activation for each cell type, we calculated firing rate change between a pre-stimulation and post-stimulation window and observed that 174 units were significantly modulated with the vast majority (91%) showing firing rate suppression. We then characterized stimulation-evoked changes in firing rate to gain insight into cell type-specific responses. Additionally, in a subset of the units, we observed that firing rate responses were modulated by stimulation distance, where local stimulation (within approximately 40 mm) could evoke instantaneous firing, whereas distant stimulation reliably suppressed firing in the same units. Finally, we analyzed a subset of units within the seizure onset zone, which exhibited unique waveform features and responses to stimulation.

This study bridges a gap in the neuromodulation field by examining the single-unit firing rate response to direct electrical stimulation of the human brain and analyzing cell-type specific firing rate responses. We show that low frequency, single-pulse stimulation broadly elicits firing rate suppression, but parameters, such as distance from the unit, can have diverse effects on firing rate responses. This work informs the neuronal basis of CCEP generation and therefore has implications for clinical mapping and informs novel ‘active probing’ strategies for precision diagnosis and neuromodulation of seizure pathophysiology in surgical cases. Moreover, this research has general implications for understanding neuromodulation via direct brain stimulation.

**Highlights:**

- Putative principal cell waveform shapes are characterized by longer trough-to-peak and full-width half max durations (s) compared to interneurons.
- Monopolar stimulation @ 3 mA generally has a widespread suppressive effect on neuronal firing, lasting approximately 1.5 s.
- Principal cells show greater suppression amplitude and longer suppression durations than interneurons.
- Stimulation within ∼ 40 mm is capable of evoking instantaneous firing, despite general suppression from stimulation at greater distances.

## Introduction

Neuromodulation with pulses of electrical current is a widespread and expanding treatment for neurological and psychiatric disorders such as Parkinson’s disease,^1^ obsessive compulsive disorder,^2^ treatment resistant depression,^3^ drug-resistant epilepsy (DRE),^4^ Tourette’s syndrome,^5^ and other neurological disorders or injury.^6^ Despite tremendous advances in the use of neuromodulation for treatment of brain disorders, its mechanism of action remains debated.^7^ Such lack of understanding of the molecular, cellular, and network mechanisms underlying neuromodulation motivates the need to understand how pulsatile electrical stimulation affects the fundamental information processing unit of the human brain: the neuron.

Direct stimulation of the human brain was first carried out intraoperatively for functional mapping.^8^ The earliest use of single-pulse electrical stimulation (SPES) to examine connectivity in the human brain was three decades ago.^9^ Since then, there has been exponential progress for the current application of neuromodulation, including examining effective connectivity of brain networks,^10,11^ identifying epileptogenic areas and functional connectivity,^12–14^, brain mapping,^15,16^, cortical connectivity,^9^ and for numerous other research applications.^17–20^ Stimulation of intracortical microelectrodes has been used more frequently to characterize single cell responses to electrical stimulation in primate^21^ and human cortices.^22–25^

Despite such long and comprehensive study of local field potential (LFP) responses to SPES (i.e., cortico-cortical evoked potentials (CCEPs)), only one previous study examined single neuron firing rate responses to SPES, likening them to patterns of activity induced by interictal epileptiform activity.^26^ In examining correlates of multi-unit firing in the LFP spectrum (i.e., broadband high-frequency or high-gamma), SPES has been shown to largely suppress these analogs of population firing^17,27–29^ however some suggest that stimulation parameters and cortical connectivity modulate firing responses.^26,30^ While there is an expansive field examining CCEPs and functional connectivity in epileptic brains,^13,17,27,31,32^ it would be valuable to know whether neuronal firing recorded from within the seizure onset zone (SOZ) in epilepsy patients express unique responses to stimulation, to inform responsive neurostimulation device use.

In this study, we sought to characterize the effect of SPES from intracortical macroelectrodes on simultaneously recorded isolated human neuron activity. To do so, we examined neuron recordings from patients with DRE while SPES was applied to patient’s stereo-electroencephalographic (SEEG) electrodes. We leveraged this unique opportunity to examine the implications of SPES on several firing rate characteristics including (1) differences in neuronal response across putative cell types, (2) precise characterization of stimulation-related changes to firing rate, (3) effects of proximity of stimulation on evoked instantaneous firing, and (4) differential response to stimulation in the SOZ. Together these analyses provide detailed characterization of human neuron responses to brain stimulation generally, and directly inform the neuronal mechanisms of neuromodulation, including treatment for epilepsy.

## Materials and methods

### Study Participants

This study was approved by the University of Utah Institution Review Board. All study participants provided informed consent prior to the implantation of Behnke-Fried (BF) microwire bundles (IRB_00114691) and prior to undergoing brain stimulation (IRB_00069440). Patients who were undergoing intracranial monitoring for DRE using SEEG electrodes consented to participate.

### Single Pulse Electrical Stimulation (SPES) Procedure

Monopolar electrical stimulation using single biphasic pulses (cathodic to anodic) were delivered every 3 to 3.5 seconds with a current intensity of 3 milliamperes and a 200-microsecond pulse width per phase (100 µs inter pulse interval; Fig. 1A). The recording and stimulation ground was a Bovie pad (Bovie Medical Corporation, Clearwater, FL) placed on the patient’s arm. A 60-second baseline period was recorded before each stimulation protocol was initiated. We stimulated each intracranial contact approximately ten times, in a randomized manner. We controlled the stimulation procedure with custom-made graphical user interface (GUI) to implement the stimulation protocol. Within this GUI, we selected minimum and maximum delivered current to 3 mA, set the number of stimulation trials for each contact to 10, and set stimulation polarity to monopolar. Stimulation trial order was set as pseudorandom, to avoid stimulating the same electrode twice in a row and to separate the next stimulated electrode by a minimum of 20 mm from the previous five stimulated electrodes,^17,27^ prohibiting overstimulation of one region of the cortex (Supplementary Fig. 1).

**Figure 1.**
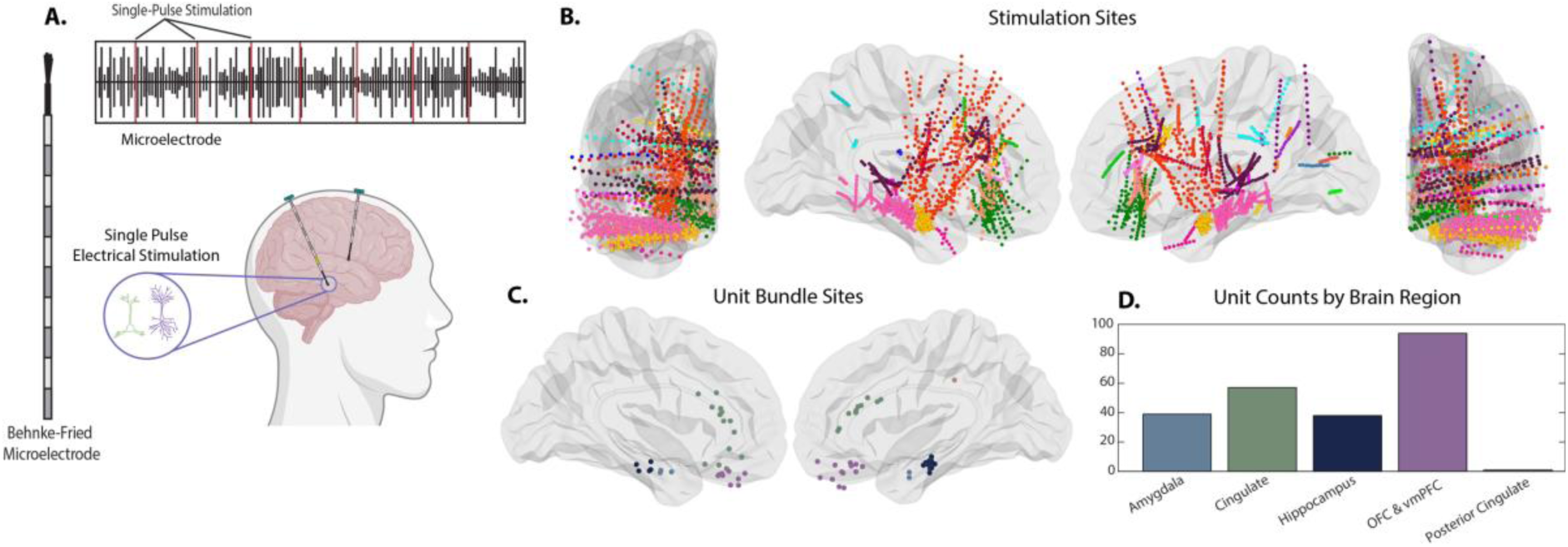
Experimental setup and electrode locations. **(A)** Single-Pulse Electrical Stimulation experiment schematic showing a cartoon of the stimulation of the selected macroelectrodes ∼10 trials each, every 3 s (top), examples of Behnke-Fried microelectrodes with bundles of 8 microwires extending from the distal tip of the electrode (left), which are inserted into the brain in order to record neuron activity (right). **(B**) Posterior and lateral views of 3D brain plot highlighting stimulation sites (N = 4329) color-coded by brain region. **(C)** Lateral view of 3D brain plot highlighting microelectrode bundle sites color-coded by brain region as shown in D. **(D)** Number of units in each of the five major regions: amygdala (light blue), cingulate (light green), hippocampus (dark blue), orbital frontal cortex & ventromedial prefrontal cortex (purple), and posterior cingulate (light brown).

### Electrodes and Recordings

SEEG electrodes have a varying number of cylindrical platinum contacts in a linear alignment. The precise location of each stimulation SEEG electrode was identified through co-registering the structural T1 MRI with a postoperative CT using the LeGUI electrode localization software (Fig. 1B).^33^ LeGUI uses Statistical Parametric Mapping software to fit patient specific anatomy to brain average-derived atlases in a probabilistic manner, from which anatomical locations are derived.^34^

Single unit activity was recorded on BF hybrid macro-micro electrodes (Ad-Tech Corp., Racine, WI), which have bundles of eight microwires, extending approximately 4 mm from the distal tip of the clinical depth electrode (Fig. 1A & C).^35^ Each participant had up to four microwire bundles implanted.^36^ BF microelectrode positions were categorized into five coarse anatomical regions: amygdala, hippocampus, OFC and vmPFC, cingulate, and posterior cingulate, defined using the Neuromorphometrics brain atlas (NMM) via the electrode localization tool LeGUI (Fig. 1D).^33^

Brain activity from BF microwires was recorded using a 128-channel data acquisition system (Blackrock Microsystems, Salt Lake City, UT) and stimulation was delivered using a 96-channel stimulation system (Cerestim, Blackrock Microsystems, Salt Lake City, UT). Electrophysiological data from microwires were sampled at 30 kHz and pseudo-differentially amplified by 10. Reference channels were selected from an intracranial depth electrode residing in white matter, for each subject (Supplementary Table 2). Recording and stimulation grounds were connected to a Bovie pad on the patient’s arm. Each recording session lasted around 40 minutes, subjects were encouraged to stay awake and relax, however there was one subject who fell asleep shortly after the recording session began (participant ID #12).

### Data Preprocessing

The neurophysiological recordings were first examined for stimulation artifact using an automatic detection algorithm. To ensure no artifacts were carried forward into neurophysiological analysis, a stimulation artifact rejection GUI in Matlab allowed for a secondary visual inspection of the data to reject remaining artifacts manually (Github: StimArtifactGUI).

### Spike Sorting and Analysis

Offline Sorter (Plexon, Dallas, TX) was used to find units within the neural data. First, the signal from each channel was set at a threshold of -3.5 RMS. BF channels were initially analyzed using an automatic spike sorting algorithm, employing a T-Distribution Expectation Maximization sorting method (parameters: degrees of freedom multiplier = 4, initial # units = 3). Then, each microelectrode channel was visually inspected for any additional units not detected by the automatic sorter.

### Classification of Putative Cell Subtypes

To classify the units into cell subtypes, we quantified features of each action potential (AP) waveform using three metrics: (1) full-width half max (FWHM), (2) trough-to-peak duration (TTP), and (3) AP asymmetry. Two clusters were observed, representing putative interneurons (INs) and principal cells (PCs). INs were classified as any unit with a FWHM threshold of <= 0.35 ms and a TTP <= 0.35 ms. To ensure this cluster selection wasn’t arbitrary and was robust, we also performed a *k*-means cluster algorithm based on the FWHM and TTP, as well as the first three principal components of the AP waveforms. Initially, we set *k* = 2, but observed a third group of units which had characteristically longer TTP durations. Therefore, we also applied *k*-means clustering with *k* = 3 (Supplementary Fig. 2). In these cases, the clearest separation in waveform feature space was between the IN and PC clusters defined by <= 0.35 ms TTP and FWHM measurements. We also clustered units based on individual subjects, brain region, significantly modulation, and direction of modulation, to confirm that no other factors contributed to the cluster pattern (Supplementary Fig. 3).

### Classification of Seizure Onset Zone Units

We also classified units as within or outside of SOZ, based on the attending epileptologist’s report. This patient-specific document provides a clinical interpretation of regions of the brain that are associated with seizure onset. Any BF microwires implanted in regions that corresponded to the clinically defined SOZ, was categorized as a SOZ unit (Supplementary Table 3).

### Neuronal Firing Rate Calculations

We calculated averaged firing rates across stimulations using the *psth* function in the chronux Matlab toolbox (chronux.org).^37^ Firing rate was convolved with an adaptive gaussian kernel starting with a 50ms bin width. For statistical comparisons for group-level and distance analysis, we standardized trial averaged firing rates, *FRz*, to control for smaller and unequal sample sizes between comparisons. To do this, we first calculated a trial by time firing rate matrix across the whole trial time window (1 s pre-stimulation to 3 s post-stimulation). We calculated the standard deviation, *SD*, of the mean firing rate, *FR*, in this time window, then created z-scored firing rate from these outcomes:

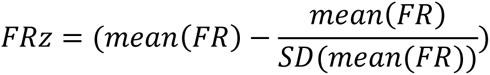

Next, we calculated took these z-scored firing rate values across time and averaged across two time windows of interest: (1) one-second before stimulation and (2) one-second after stimulation. We conducted a *t*-test over trials to see whether the standardized firing rates were significantly different from pre to post stimulation. This determined whether the unit was significantly modulated by the stimulation. A significant positive change defined firing rate enhancement, whereas a significant negative change defined firing rate suppression.

### Neuronal Feature Definitions

We sought to precisely characterize features of the firing rate response to SPES by expanding upon some previously reported metrics:^26^

*Baseline Frequency (Hz):* The trial averaged firing rate during a one second pre-stimulation time window (-1.1 s – -0.1 s).

*Post-Stimulation Firing Rate (Hz):* The trial average firing rate during a one second post-stimulation time window (0.1 s – 1.1 s).

*Suppression Amplitude (Hz):* The lowest instantaneous trial averaged firing rate during suppression during a one second post-stimulation time window (0.01 s – 1.11 s).

*Suppression Latency (ms):* Time taken from stimulation at time zero to the timepoint that the minimum suppression amplitude occurs.

*Duration of the Suppression (ms):* Difference in time since the instantaneous trial averaged firing rate crossed downwards and upwards 2 SD lower threshold below the baseline frequency.

*Area Under the Curve (AUC)*: We calculated AUC to observe potential differences between cell types. For this analysis, we inverted standardized firing rates, then did numerical integration on the data greater than zero to calculate an AUC frequency for firing rate under the suppression threshold.

### Distance Versus Firing Rate Change Point Analysis

To calculate the Euclidean distance at which firing rate responses transitioned from evoked to suppressed, we fit sigmoidal curves to mean firing rates across stimulations from each stimulation electrode, sorted by Euclidean distance from each microwire bundle. The Euclidean distance between each electrode pair is defined by the length of the shortest line segment connecting the two electrode locations in three-dimensional space. Accordingly, we calculated the three-dimensional Euclidean distance, 𝑑_𝑖_, between the location of the microelectrode bundle, (𝑥_𝑢_, 𝑦_𝑢_, 𝑧_𝑢_) and the location of each stimulating electrode, (𝑥_𝑖_, 𝑦_𝑖_, 𝑧_𝑖_),

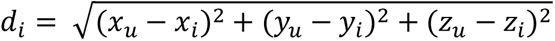

Then, using the Matlab function *sigmd_fit*, we fit a sigmoid to each unit’s mean firing rate between 50 and 300 milliseconds following stimulation, sorted by Euclidean distance. This procedure yielded a model of firing rate versus stimulation distance. We then operationally defined the change point, or Euclidean distance at which firing rate changed from enhancement to suppression, as the x50 parameter, or inflection point, of the sigmoidal curve. This procedure is depicted in Supplementary Fig. 4.

### Statistical Tests

All statistical results are reported in the text as mean ± standard deviation.

*Proportion Tests.* Chi-square proportion tests (alpha level = 0.05) were used to test whether (1) more INs were suppressed than PCs (2) more INs were enhanced than PCs (3) more SOZ units were suppressed than non-SOZ units (4) more SOZ units were primarily INs versus PCs, and (5) more SOZ INs were suppressed than SOZ PCs.

*Rank-sum Tests.* Two-tailed Wilcoxon rank-sum tests (alpha level = 0.05) were used to test for numerous group-level differences: (1) differences between cell type waveform metrics (TTP, FWHM, and asymmetry), (2) differences between non-SOZ and SOZ unit waveform metrics (TTP, FWHM, and asymmetry), (3) average and z-scored FRs between INs and PCs, and between SOZ and non-SOZ units, for (a) baseline frequency, (b) post-stimulation frequency, (c) suppression amplitude, (d) suppression latency, (e) suppression duration, (f) suppression thresholds, and (h) area under the curve.

*Regression Analysis.* To examine whether there were any associations between neuronal features across neurons we fit linear regression models for (1) suppression amplitude and suppression latency, (2) suppression amplitude and suppression duration, and (3) suppression duration and suppression latency. These models were fit for all units and between cell types and SOZ and non-SOZ units.

## Results

### Patients and Data

We applied low frequency monopolar SPES to the majority of SEEG contact in 30 participants (16 female, 36 ± 10.39 years of age; Fig. 1A & Supplementary Table 1). Across all participants, stimulation was applied to a total of 4329 sEEG contacts after 3.03% of stimulation channels were rejected during data preprocessing (M ± SD = 148 ± 38.8 contacts per participant; Fig. 1B). In total, 228 isolated units were identified in recordings from 76 microwires (2.53 ± 0.63 microelectrode bundles implanted per patient; Supplementary Table 2; Fig. 1C & D) averaging approximately three units recorded per microelectrode bundle.

### Cell Types Have Distinct Waveform Metrics

Out of the 228 total identified units, 191 (83.8%) were classified as PCs and 37 (16.2%) were classified as INs (Fig. 2A-C). These units were sampled from five key anatomical regions amygdala (6 IN, 33 PC), OFC & vmPFC (13 IN, 81 PC), hippocampus (9 IN, 29 PC), cingulate (9 IN, 48 PC), and posterior cingulate (0 IN, 1 PC; Fig. 2D).

**Figure 2.**
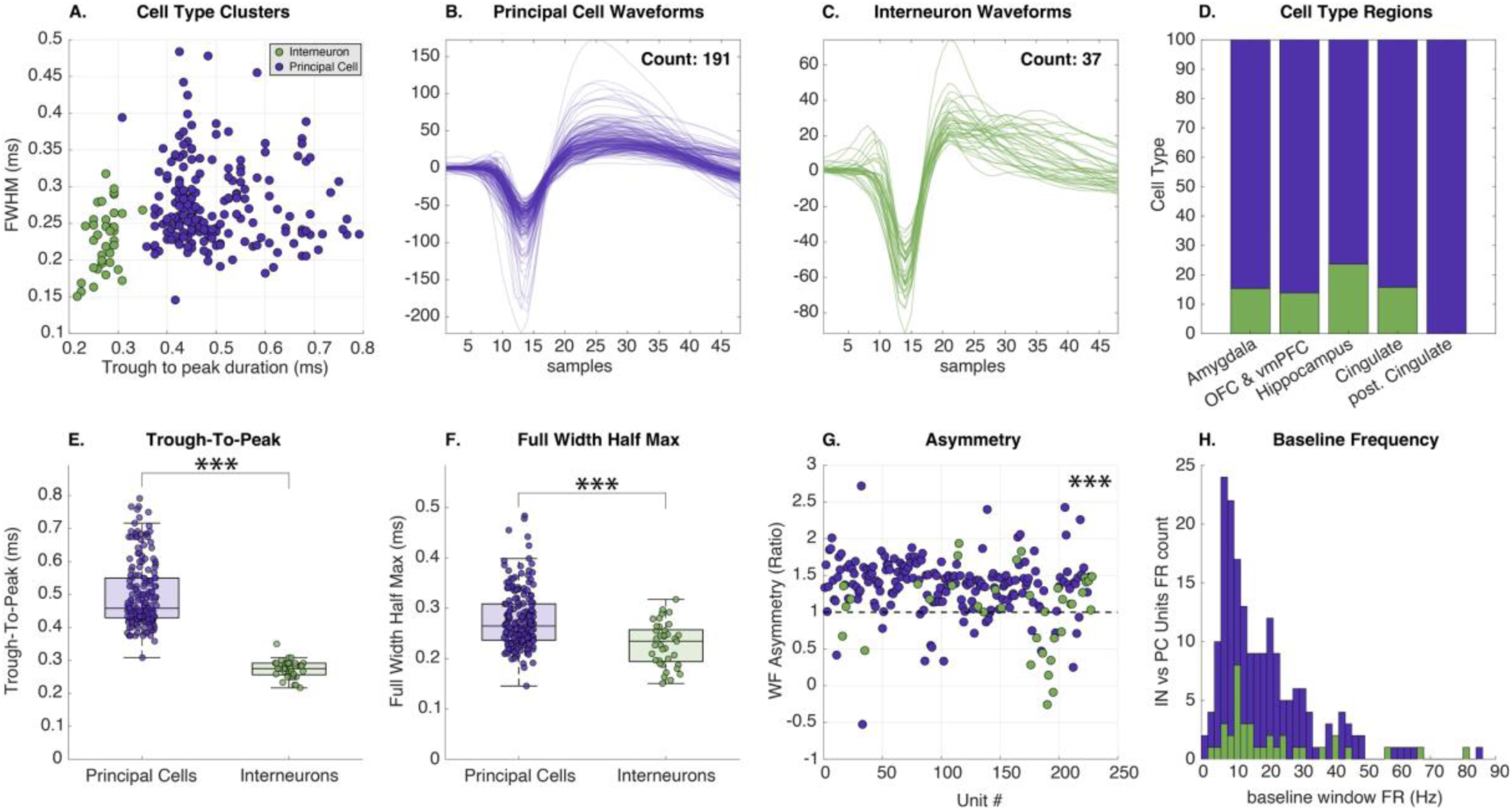
Cell Type Waveform Classification. **(A)** a scatter plot of waveform features for each average waveform, color coded by cell type (green: INs, purple: PCs). **(B)** Putative PC waveforms (purple cluster in panel A). **(C)** Putative IN waveforms (green cluster in panel A). (**D**) Cell type proportions for the five main regions: amygdala (15.4% INs), OFC & vmPFC (13.8% INs), hippocampus (23.68% INs), cingulate (15.79% INs), and posterior cingulate (0% INs). **(E)** TTP (ms) metric for PCs (0.5 ms ± 0.1) and INs (0.27 ms ± 0.03). **(F)** FWHM (ms) metric for PCs (0.28 ms ± 0.06) and INs (0.23 ms ± 0.04). **(G)** Asymmetry (Ratio) for PCs (1.4 ± 0.4) and INs (1.04 ± 0.5). **(H)** Baseline Frequency (Hz) for PCs (21.63 Hz ± 15.34) and INs (22.2 Hz ± 18.6). [Significance (*p* < 0.05*, 0.01**, 0.001***]

The three main metrics used to quantify the waveforms were TTP duration (ms), FWHM duration (ms) and asymmetry (Fig. 2E-G). There was a significant difference between TTP for INs (0.27 ms ± 0.03) and PCs (0.5 ms ± 0.1; *rank-sum* = 24720, *z* = 9.61, *p* < 0.001), FWHM for Ins (0.23 ms ± 0.04) and PCs (0.28 ms ± 0.06; *rank-sum* = 22892, *z* = 4.55, *p* < 0.001), and asymmetry for INs (1.04 ± 0.52) and PCs (1.38 ± 0.38; *rank-sum* = 22731, *z* = 4.12, *p* < 0.001). Our waveform classification threshold therefore robustly highlighted distinct waveform features for different cell types.

### Stimulation Causes Widespread Firing Rate Suppression

We sought to understand whether cell types had disparate responses to stimulation by testing for differences in firing rate response features between cell types. Overall, 174 (76.3%) units out of the 228 total recorded units significantly responded to stimulation. From the responsive sample of units (examples: Fig. 3A & B), 159 (91.4%) showed firing rate suppression post-stimulation (Fig. 3C & Supplementary Fig. 5) and 15 (8.6%) showed firing rate enhancement post-stimulation (Fig. 3D). Enhanced units had a baseline frequency of 7.3 ± 3.3 Hz, with a post-stimulation maximum frequency of 9.9 ± 5.1 Hz. Between cell types, enhanced INs had a baseline frequency of 8.3 ± 3.9 Hz and a maximum post-stimulation firing rate of 8.5 ± 4.7 Hz, while enhanced PCs had a baseline frequency of 7.1 ± 3.4 Hz and a maximum post-stimulation firing rate of 10.1 ± 5.3 Hz. Due to low sample sizes, we were not powered to statistical test for differences in firing rate response features among enhanced units.

**Figure 3.**
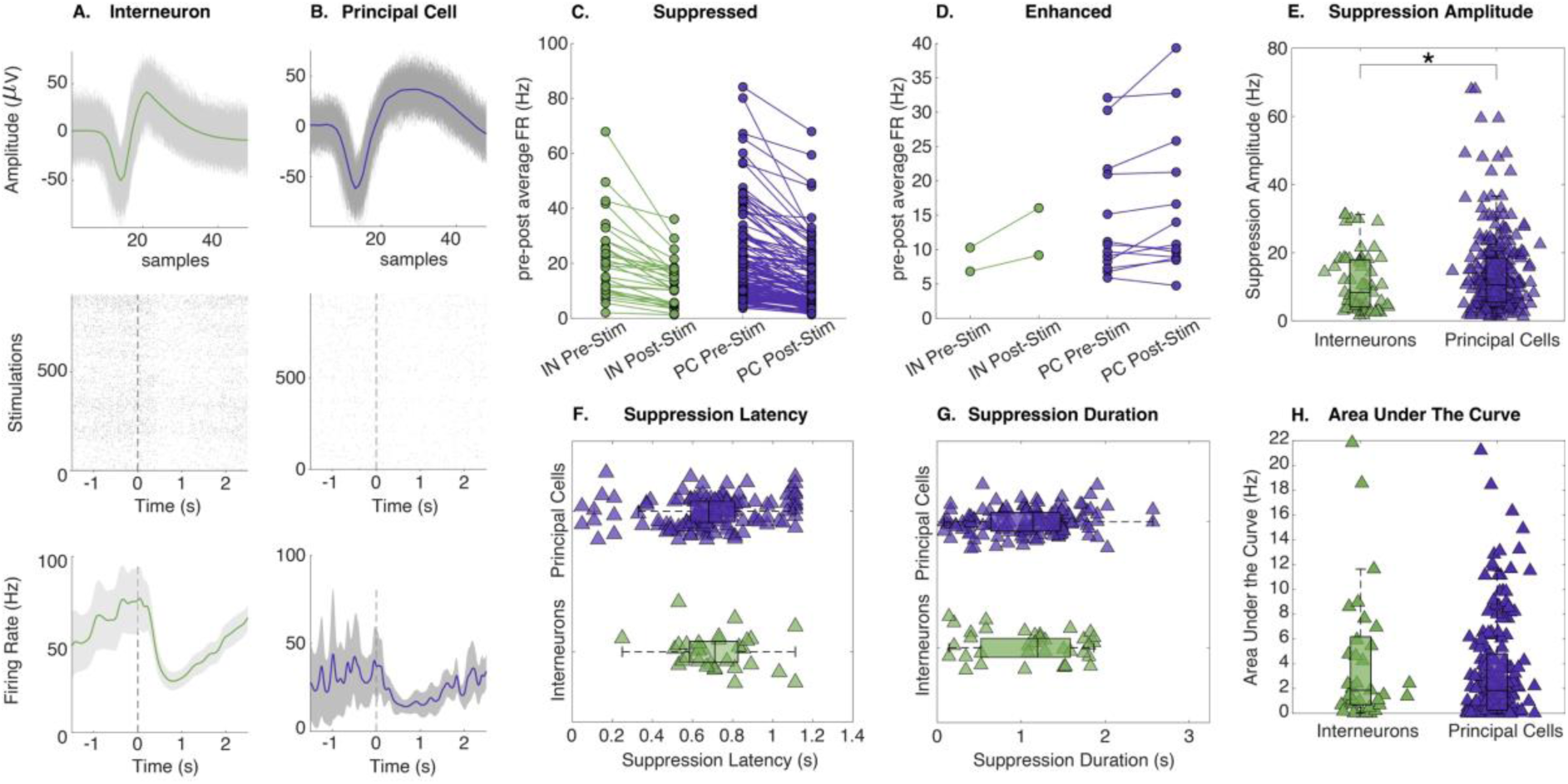
Cell Types Neuronal Responses. **(A)** Example individual IN waveform from left orbital frontal cortex (green) with SEM patch (grey) *(top)*, spike raster plot *(middle)*, average firing rate (green) with SEM patch (grey) *(bottom)*. **(B)** Example individual PC waveform from left orbital frontal cortex (purple) with SEM patch (grey) *(top)*, spike raster plot *(middle)*, average firing rate (purple) with SEM patch (grey) *(bottom)*. Dashed lines in firing rate and raster plots represent stimulation at time zero. **(C)** Mean changes in average firing rates for suppressed INs (green) and PCs (purple). **(D)** Mean changes in average firing rates for enhanced INs and PCs change in average firing rates. **(E)** Box plot showing median suppression amplitude and interquartile range for each cell type with an underlaid scatter plot showing mean suppression amplitude for each unit. **(F)** Box plot showing median suppression latency and interquartile range for each cell type with an underlaid scatter plot showing mean suppression latency for each unit. **(G)** Box plot showing median suppression duration and interquartile range for each cell type with an underlaid scatter plot showing mean suppression duration for each unit. **(H)** Box plot showing median area under the curve and interquartile range for each cell type with an underlaid scatter plot showing mean area under the curve for each unit. [Significance (*p* < 0.05*, 0.01**, 0.001***]

Since most recorded units exhibited firing rate suppression, we went on to characterize features of this response in the subset of significantly suppressed units. Suppressed units exhibited a mean baseline firing rate of 21.8 ± 16.0 Hz which was suppressed by stimulation to a minimum firing rate of 13.2 ± 11.0 Hz. Suppression latency was 0.69 ± 0.24 s, with suppression lasting 1.08 ± 0.55 s after crossing a suppression threshold of 14.3 ± 14.1 Hz. Proportion tests revealed no significant differences in quantities of suppressed cell types (INs = 31/33, PCs = 128/141, χ^2^ = 0.34, *p* = 0.56).

We tested for across-unit associations between suppression duration, amplitude, and latency, by regressing these response features against each other. Across all units, we found a significant association between suppression amplitude and suppression duration (*t*(116) = 2.35, *p =* 0.0202), but not between suppression amplitude and latency (*t*(116) = -1.49, *p =* 0.1382) or suppression duration and latency (*t*(116) = -1.408, *p =* 0.162). SPES therefore caused widespread firing rate suppression among both cell types.

### Cell Types Exhibit Different Firing Rate Features in Response to Stimulation

Having found that cells were overall suppressed by SPES, we next tested for differences in each firing rate response feature between cell types. Suppressed INs exhibited a mean baseline firing rate of 22.2 ± 18.6 Hz, which stimulation suppressed by a mean of 11.1 Hz (∼50.0% firing rate decrease) to a minimum firing rate of 11.1 ± 8.7 Hz, while suppressed PCs exhibited a similar mean baseline firing rate of 21.6 ± 15.3 Hz, which stimulation suppressed by a mean of 7.9 Hz (∼36.6%) to a minimum firing rate of 13.7 ± 11.5 Hz. When comparing cell type responses, INs had higher baseline firing rates (IN = -3.23, PC = -4.0; *z* = 2.64, rank-sum = 3088, *p* = 0.0083) (Fig. 2H) and post-stimulation firing rates (IN = -3.2, PC = -3.9; *z* = 2.47, rank-sum = 3048, *p* = 0.014), but PCs had greater magnitude of suppression (IN = -3.8, PC = -4.6; *z* =2.32, rank-sum = 3013, *p* = 0.021; Fig. 3C-E).

Suppression latency for INs was 0.72 ± 0.19 s, with suppression lasting for an average of 1.1 ± 0.6 s after crossing the suppression threshold of 13.7 ± 14.5 Hz (Fig. 3E-G). Suppression latency for PCs was 0.69 ± 0.25 s for PCs, with suppression lasted for an average of 1.08 ± 0.55 s, after crossing the suppression threshold of 14.5 + 14.3 Hz (Fig. 3E-G). Between cell types, no differences were observed for suppression latency, suppression duration, suppression threshold, or AUC (Fig. 3H). Despite differences in average firing rates, PCs were suppressed to a greater degree than INs, yet both cell types demonstrated similar suppression latencies and durations. Still, we were interested in testing whether there were any associations between suppression features across cells. We found a significant positive association between suppression amplitude and duration of PCs (*t*(90) = 2.018, *p =* 0.047), and did not see the same effect for INs (*p* > 0.05). These findings emphasize PC-specific increases in suppression from stimulation.

### Stimulation From Local Sites Can Cause Evoked Firing

Despite the general firing rate suppression reported above, we observed stimulation evoked instantaneous firing from several units, and that these responses were elicited from stimulation nearest to the microelectrodes. We thus sought to quantify how stimulation distance modulated these firing rates. To do this, we sorted firing rates from each stimulation electrode by its Euclidean distance from each unit and found that 40 (17.5%) units showed evoked firing rates from stimulation on SEEG contacts (Fig. 4A-D). Of those 40 units, 31 showed evoked firing in response to stimulation near the microelectrodes and 9 showed evoked firing in response to stimulation further away from the electrode. To characterize the abrupt changes between evoked and suppressed firing that we noticed in these units, we fit sigmoidal curves to the mean firing rates across simulating electrode distances to model their inflection points. The inflection points of these sigmoids occurred at an average distance of 39.1 ± 14.0 mm from the microelectrode bundle (Fig. 4E). From the 31 local responsive units, the average firing rate below the distance change point threshold for evoked firing was 1.8 Hz ± 0.8 and above the distance change points threshold was 1.15 Hz ± 0.42. On average, this evoked firing lasted for approximately 200 ms post-stimulation before the firing rate became characteristically suppressed (Fig. 4A & B).

**Figure 4.**
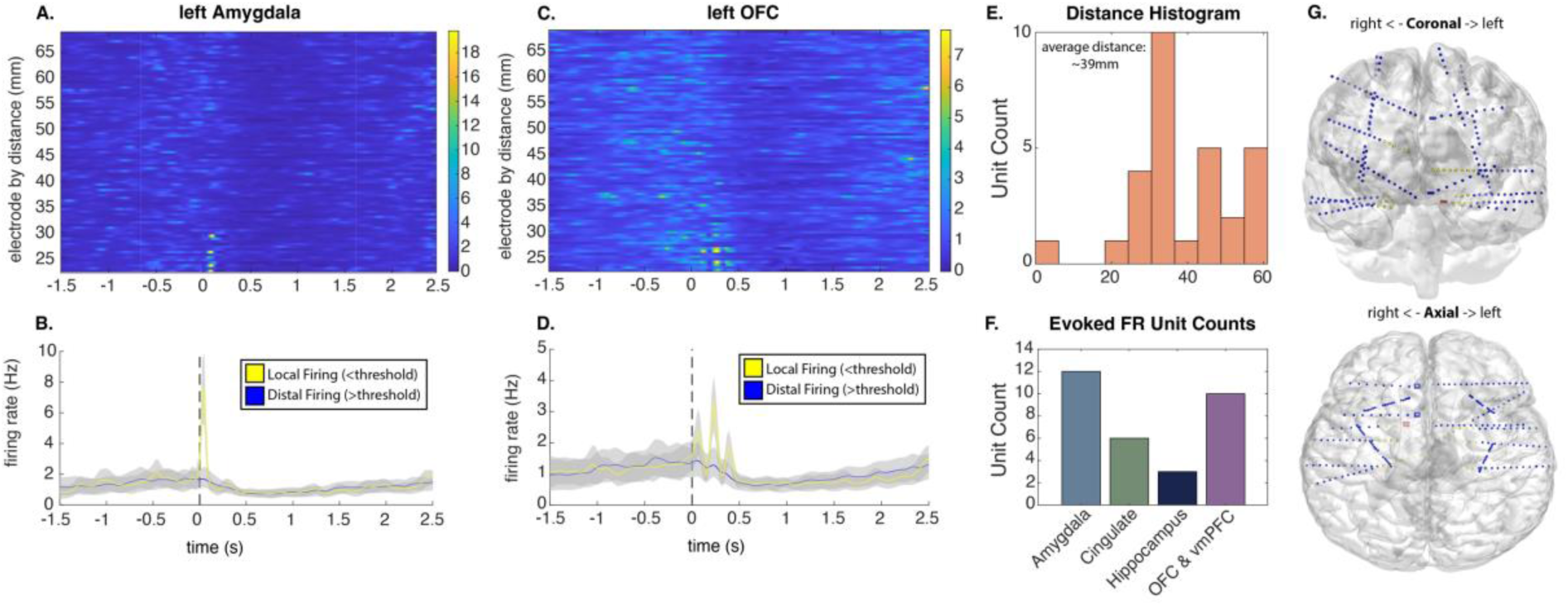
Evoked Firing Rate by Euclidean Distance. **(A)** Example in left amygdala of unit firing rate ordered by Euclidean distance of stimulation. **(B)** Firing rate of left amygdala unit below evoked firing threshold (<=34mm; yellow) and above (> 34mm; blue) around time of stimulation (time 0s). **(C)** Example in left Orbital Frontal Cortex of unit firing rate ordered by Euclidean distance of stimulation. **(D)** Firing rate of left orbital Frontal cortex unit below evoked firing threshold (<28mm; yellow) and above (>= 28mm; blue) around time of stimulation (time 0s). **(E)** Distance histogram of the firing rate roll off for evoked near unit firing (mean = ∼39mm). **(F)** Regional summary of unit counts with evoked near firing caused by stimulation (amygdala = 12, hippocampus = 3, cingulate = 6, orbital frontal cortex = 10, and posterior cingulate (not plotted) = 0). **(G)** Example of an amygdala unit and evoked firing from all stimulation sites. (*top*) coronal view brain, (*bottom*) axial view of brain. Colormap highlights stimulation sites firing rates (Hz).

Next, we examined whether this firing rate modulation was related to the cell type that the unit was previously classified as. Overall, 32 of the 40 units were principal cells (24 near evoked firing rate) and 8 were interneurons (7 near evoked firing rates). We ran a proportion test to examine whether cell type modulated evoked firing closer to the unit, however this was confirmed insignificant (χ^2^ = 0.574, *p* = 0.449). Finally, we examined whether there were specific brain regions from which stimulation was associated with increases in evoked firing. Proportion tests revealed that amygdala units were significantly more likely to show evoked firing rate responses to stimulation compared to OFC (χ^2^ =12.313, *p* = 4.4990e-04), hippocampus (χ^2^ = 10.798, p = 0.001), and cingulate units (χ^2^ = 10.111, *p* = 0.0015). No other regions had significant proportions of evoked firing units.

### SOZ PCs Exhibit Longer Trough-To-Peak Durations

We recorded 27 units that were clinically defined as within the SOZ (Fig. 5A-C). These units were sampled across our coarse anatomical regions with the highest proportion of SOZ units from hippocampus (29.0%) (SOZ unit count: amygdala = 4; OFC & vmPFC = 6; hippocampus = 11; cingulate = 6; Fig. 5). Proportion tests revealed no significant difference between the number of SOZ units that were classified as INs versus PCs (χ^2^ = 0.118, *p* = 0.731).

**Figure 5.**
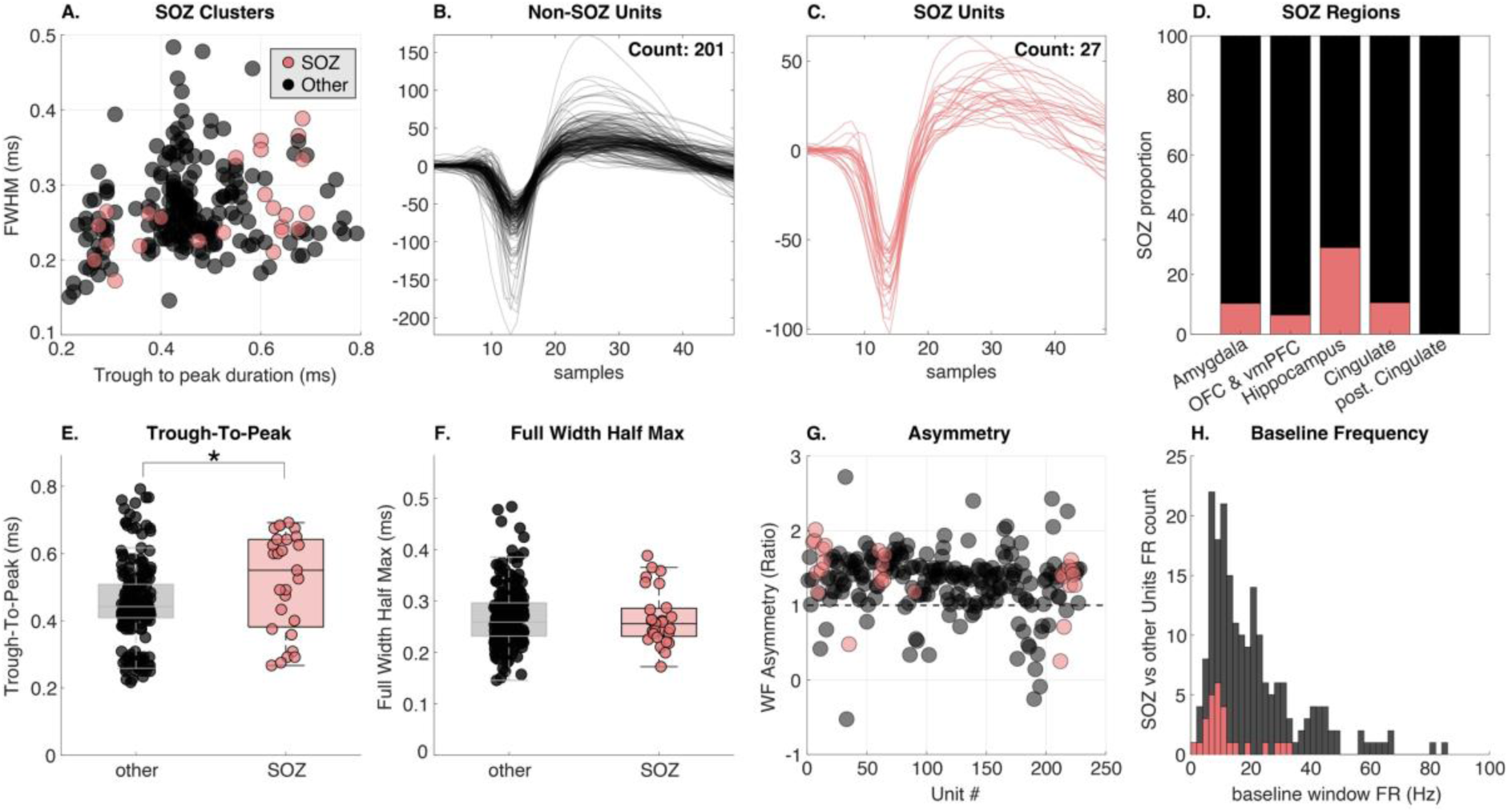
SOZ Waveform Classification. **(A)** a scatter plot of waveform features for each average waveform, color coded by cell type (pink: SOZs, black: non-SOZs). **(B)** Non-SOZ waveforms (black cluster in panel A). **(C)** SOZ waveforms (pink cluster in panel A). **(D)** proportions for the five main regions: amygdala (10.3% SOZ), OFC & vmPFC (6.4% SOZs), hippocampus (29.0% SOZs), cingulate (10.5% SOZs), and posterior cingulate (0% SOZs). (E) TTP duration for non-SOZ (0.45 ± 0.12 ms) and SOZ cells (0.52 ± 0.15 ms). **(F)** FWHM duration for non-SOZ (0.27 ± 0.06 ms) and SOZ cells (0.27 ± 0.06 ms). **(G)** Asymmetry (Ratio) for non-SOZ (1.3 ± 0.4) and SOZ units (1.4 ± 0.4). **(H)** Baseline Frequency (Hz) (- 1.1s -0.1s before stimulation) for non-SOZ (23.0 ± 16.3 Hz) and SOZ units (11.9 ± 8.5 Hz). [Significance (*p* < 0.05*, 0.01**, 0.001***]

We observed a significant increase in TTP duration for SOZ units (0.52 ± 0.15 ms) compared to non-SOZ units (0.45 ± 0.12 ms; *z* = 2.321, *rank-sum* = 3787, *p* = 0.0203; Fig. 5E). However, we saw no significant differences between FWHM (SOZ = 0.27 ± 0.06 ms; non-SOZ = 0.27 ± 0.06 ms; *z* = -0.345, *rank-sum* = 2971, *p* = 0.73; Fig. 5F), or asymmetry (SOZ = 1.4 ± 0.4; non-SOZ = 1.3 ± 0.4; *z* = 1.603, *rank-sum* = 3560, *p* = 0.109; Fig. 5G). Since this increased TTP duration appeared to occur in PCs, and the majority of our SOZ units were classified as PCs (81.5%), we recapitulated this analysis with only PC units to examine potential cell type-specific properties that SOZ INs may be diluting. Again, we observed a significant increase in TTP duration for SOZ units (0.57 ± 0.11 ms) compared to non-SOZ units (0.49 ± 0.09; *z* = 3.195, *rank-sum* = 2845, *p* = 0.0014), but no difference in FWHM (*p* = 0.975) or asymmetry (*p* = 0.08). There was no difference in baseline frequency for SOZ or other units (*p* > 0.05). These results show that SOZ PCs have wider overall AP waveforms.

### SOZ Neuron Responses to Stimulation

Finally, we wanted to see if the subset of SOZ units responded differently to stimulation, compared to units outside of the SOZ (Fig. 6A & B). Among the 27 SOZ units, 18 were suppressed, 3 were enhanced and 6 had no response to stimulation (Fig. 6C & D). Overall, SOZ units exhibited shorter suppression durations (*z* = -2.044, *rank-sum* = 383, *p* = 0.041) and smaller suppression threshold (*z* = -3.525, *rank-sum* = 791, *p* < 0.0001) compared to non-SOZ units (Fig. 6G). We found no significant differences for the other firing rate response features (*p*’s > 0.05) (Fig. 6E, F, & H). As we saw differences in firing rate features across cell types, we looked within cell type at suppressed INs (SOZ = 5; non-SOZ = 26) and PCs (SOZ = 13; non-SOZ = 115). However, proportion tests revealed no differences in the proportion of SOZ units that were suppressed, between cell types (χ^2^ = 0.889, *p* = 0.346). These results detail diminished responses to SPES in SOZ units.

**Figure 6.**
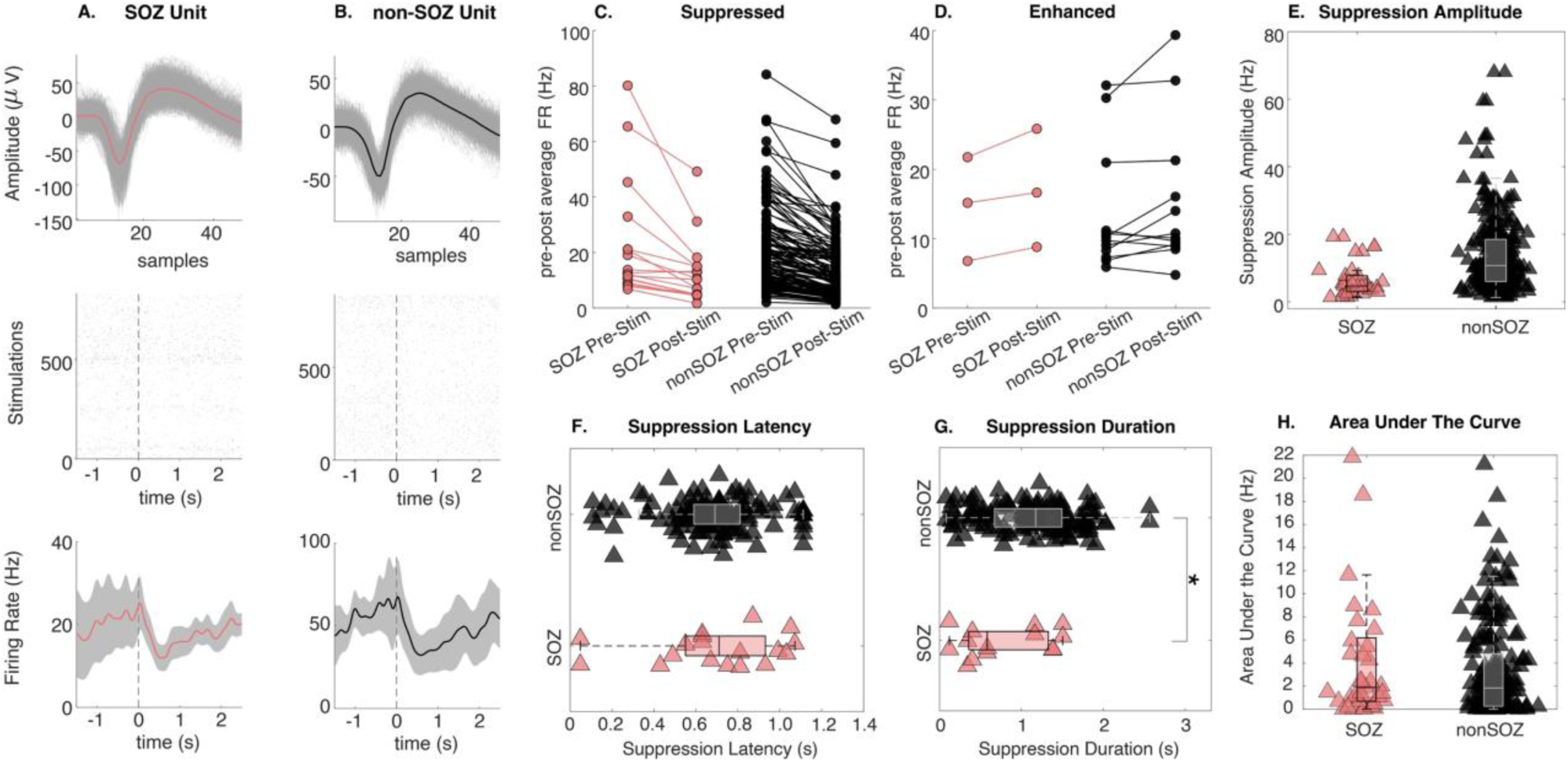
SOZ Neuronal Responses. **(A)** Example individual SOZ waveform from right anterior cingulate cortex (pink) with SEM patch (grey) *(top)*, spike raster plot *(middle)*, average firing rate (pink) with SEM patch (grey) *(bottom)*. **(B)** Example individual non-SOZ waveform from left amygdala (black) with SEM patch (grey) *(top)*, spike raster plot *(middle)*, average firing rate (black) with SEM patch (grey) *(bottom)*. Dashed lines in firing rate and raster plots represent stimulation at time zero. **(C)** Mean changes in average firing rates for suppressed SOZs (pink) and non-SOZs (black) **(D)** Mean changes in average firing rates for enhanced SOZs (pink) and non-SOZs (black) **(E)** Box plot showing median suppression amplitude and interquartile range for each cell type with an underlaid scatter plot showing mean suppression amplitude for each unit. **(F)** Box plot showing median suppression latency and interquartile range for each cell type with an underlaid scatter plot showing mean suppression latency for each unit. **(G)** Box plot showing median suppression duration and interquartile range for each cell type with an underlaid scatter plot showing mean suppression duration for each unit. **(H)** Scatter plot showing area under the curve calculation of suppression between SOZs and non-SOZs. [Significance (*p* < 0.05*, 0.01**, 0.001***]

Suppressed SOZ units exhibited a mean baseline firing rate of 11.965 ± 8.495 Hz, which stimulation suppressed by a mean of 5.5 Hz (∼45%) to a minimum firing rate of 6.5 ± 5.2 Hz. Suppression latency was 0.69 ± 0.30 s, with suppression lasting for an average of 0.79 ± 0.56 s, after crossing the suppression threshold of 5.6 ± 7.7 Hz.

Suppressed non-SOZ units exhibited a mean baseline firing rate of 23.0 ± 16.3 Hz, which stimulation suppressed by a mean of 9 Hz (∼39%) to a minimum firing rate of 14.1 ± 11.3 Hz. Suppression latency was 0.69 ± 0.23 s, with suppression lasting for an average of 1.10 ± 0.55 s, after crossing the suppression threshold of 15.4 ± 14.6 Hz. Finally, we found significant positive associations between suppression amplitude and suppression durations (*t*(106) = 2.041, *p =* 0.044), and duration and latency (*t*(106) = -2.188, *p =* 0.031), in non-SOZ firing rate responses. No significant effects were observed for SOZ units (*p*’s > 0.05), however there appears to be bi- directional associations between duration and latency of non-SOZ units and SOZ units (Supplementary Fig. 6).

## Discussion

Neuromodulation is rapidly progressing field with numerous applications. Despite the widespread use of intracranial stimulation for studying, diagnosing, and treating both neurological and psychiatric disorders, *very little is known about how human neurons fundamentally respond to brain stimulation*. In this study, we have begun to close this knowledge gap by characterizing isolated neuron responses to single pulses of intracranial stimulation. Additionally, we studied specific classes of neurons, using sophisticated waveform metric analysis to isolate putative inhibitory INs from excitatory PCs, from which we examined cell-specific responses to stimulation. We observed numerous differences in putative cell type specific responses to stimulation; while INs had higher baseline firing, PCs showed greater overall magnitude of suppression and specific associations between suppression magnitude and duration. Moreover, we showed exciting findings regarding distance-modulated stimulation, such that stimulation within ∼40 mm can evoke instantaneous firing. Finally, we highlight intriguing results suggesting single units within clinically defined SOZ have wider waveform shapes and less suppression in response to stimulation. These findings encapsulate detailed characterization and quantification of human neuron responses to brain stimulation, and directly inform the neuronal mechanisms of neuromodulation, including treatment for epilepsy.

### Single Pulse Electrical Stimulation Mainly Causes Suppression of Human Neurons

Overwhelmingly, we observe suppression of single unit firing rate in response to single pulse stimulation. Typically, SPES suppresses firing rate by approximately 12 Hz and lasts approximately 1.5 s, before stabilizing to pre-stimulation firing rates. Our findings show that the research field’s push for higher amperage and bipolar stimulation to elicit cortical responses to stimulation is not always necessary, as we observe that low-level, monopolar stimulation induces neuronal suppression. A very small subset of neurons responded to stimulation by enhancing their firing rate after stimulation. These findings support previous work that found delayed high frequency suppression has also been observed after stimulation.^27^ The only previous research into the effects of macroelectrode SPES on unit firing, likened changes in firing rate to those induced by interictal epileptiform discharges.^26,38^ We extend these findings by more precisely characterizing specific cell-type differences to SPES across a larger population of cells.

### Characteristics of Action Potential Waveforms

One such use of microelectrode recordings is for the classification of putative cell types, such as PCs^39^ and GABAergic INs.^40^ Pioneering work on the classification of these neuronal subtypes has informed mechanistic and computational features of units.^41^ For instance, units that enhance other units at monosynaptic, short (< 3 ms) latency were considered excitatory pyramidal cells and units that suppress discharges were classified as inhibitory INs.^41^ The interactions between different cell types modulate complex cortical dynamics and neural networks^39,42^ and so classifying these responses to stimulation gives us new insights into human brain networks and the connections between them. Previous work shows that these neuronal subtypes can be reliably classified by extracellular features (i.e., firing rate and spike patterns),^43^ more recently, a growing standard is the use of the FWHM of a waveform to determine specific cell types.^44^ We used the FWHM alongside other waveform metrics to quantify differences between our recorded cell types;^43,45^ around 16% of the sampled units were classified as IN. We found INs have shorter FWHM and TTP durations compared to PCs as well as greater asymmetry. Our findings support previous research starting with a comprehensive review of NHP and animal model stimulation studies which found an average FWHM of ∼0.36 ms for INs.^46^ As well as animal studies, similar FWHM for INs have been found in human studies using microelectrode recordings (see Supplementary Fig. 3 in cited paper).^47^ Finally, our metric thresholds are not dissimilar (within 10 ms) to guidance on TTP cutoff for putative cell-type classification.^48^ Thus, we are confident that our reported thresholds represent general properties of AP waveforms.

### Single Pulse Electrical Stimulation Highlights Cell-Type Specific Responses

A major gap in neuromodulation research is how single units respond to neocortical stimulation. We tackle this problem by analyzing new features and looking at specific cell-type responses.^26^ We highlight unique responses of cell-types in response to stimulation. We found INs had higher baseline firing rates and post-stimulation evoked firing, supporting the overarching cellular literature that inhibitory GABAergic INs have higher firing rates than excitatory glutamate PCs.^49^ After stimulation, we saw PCs exhibited greater magnitudes of suppression compared to INs. We suspect the suppression differences we observed between cell types was related to firing rate recovery and the cortical network dynamics between excitatory and inhibitory neurons.^50^ Previous studies examining seizure propagation have shown that excitatory PCs and inhibitory INs have different rebound effects post-inhibition, with a slower rebound of post-inhibitory excitatory neurons.^51–54^ It is likely that stimulation dampens firing of PCs to a greater extent than INs, but as INs have higher firing rates in general, the rebound is relatively similar. This is supported by our finding that no cell type differences were observed between suppression durations, suggesting this is a property of the cortical roles and dynamics between inhibitory INs and excitatory PCs.^49,50^ As previous research has shown IEDs and SPES to elicit similar responses,^26,38^ we expect SPES is eliciting a similar response in excitatory PCs, post-stimulation. Furthermore, we saw PC-specific associations between the magnitude of and duration of suppression across cells, further highlighting these features of PCs.

### Neurons with High Firing Rates are Modulated by Distance

This work reveals another useful application of SPES: to quantify neuronal firing rate responses, relative to distance from the stimulation site. Previous work assessed evoked CCEP signal responses in distant or nearby cortical regions found that local (<15 mm) grey white boundaries and distant (>15 mm) white matter boundaries elicited the greatest CCEP responses to stimulation.^55^ Moreover distant CCEP networks have shown to be most similar to diffusion tensor imaging (DTI) networks, when used to measure passive (resting state) and active (SPES) connectivity in the human brain.^56^ Finally, one study found observing an initial high frequency firing rate within the first 100 ms post-stimulation for 25% of local neurons (within 3cm of stimulation electrode).^26^ Our findings support previous work as we observed a subset of units that had stimulation distance specific responses, whereby close stimulation site, within 40 mm, evoked an increased firing rate from the neuron for around 200 ms (Fig. 4).^17,27,57^

### Disparate APs for Seizure Onset Zone Cells

As the participant cohort are recruited from DRE patients, we often record from clinically relevant SOZ sites. Previous epilepsy research suggests that SOZ waveforms show a reduction in amplitude and increase in FWHM^51^ and TTP^44^ durations *during* seizures. While we did not record during seizures, our stimulation likely recapitulates activity that is similar to interictal activity. We did not observe a change in FWHM, but interestingly, found that cells within SOZ had characteristically and consistently longer TTP durations, by approximately 60ms. We speculate that this waveform shape reflects neurobiological deviations in ionic stability, related to the impact of ictal activity. Physiologically, the TTP timeframe reflects repolarization and refractory periods of an AP, when potassium (K+) ions leave the cell membrane through voltage-gated channels, restoring the resting membrane ionic potential (∼ -70mV) after briefly overshooting (refractory period). Our findings suggest that cell waveforms within seizure-affected regions of the brain have longer TTP durations, likely due to cellular dysregulation and imbalance of ion content (i.e., slower repolarization). Moreover, we find that SOZ units have smaller suppression magnitudes in response to stimulation, which may also be reflective of aberrant connectivity or sensitivity characteristic of the SOZ. The broad suppression we observe could contribute mechanistically to the recent observations that stimulation in low-risk periods correlates with seizure reduction from RNS,^58^ or that low frequency stimulation is capable of suppressing seizures.^59^

## Limitations

One fundamental limitation of any neurostimulation study, is that the parameter space for neurostimulation is enormous. Our study is not immune from such myopia, as we only examined 3 mA pulses with 200us pulse width per phase.

We define the seizure onset zone based on clinical epileptologist’s interpretation from long-term monitoring EEG recording. This SOZ selection method has some drawbacks in that its based solely on low frequency LFP and visual review of an attending epileptologist. An alternative strategy would be to compare these brain regions to surgical outcomes, to function measures of seizure spread such as the Epileptogenicity Index’ (EI),^60,61^ ictal phase-locked high gamma,^62^ or the doppler method.^63^ Furthermore, we have limited numbers of SOZ units (n = 27, 11%) so we encourage readers to interpret these findings with caution. We will continue to collect SOZ units as electrode placement permits and see how our SOZ-related findings persist with increased statistical power.

## Summary

The potential of SPES as a neuromodulation technique has been limited to the niche clinical research areas mentioned above and more specifically confined to LFP analysis,^64^ instead of single-unit firing rate responses. We show that low-voltage, single-pulse electrical stimulation is enough to evoke specific suppression responses by inhibiting neuronal firing rates. This study is the first to characterize single units, including units within SOZ, and classify putative cell types in response to stimulation, highlighting cell type-specific responses in the human brain. These findings have clear clinical implications for patients with neurostimulation devices, such as RNS, such that it can help inform the anatomical placement of such devices. These findings provide important context for human neurostimulation studies and inform the neuronal mechanisms of therapies involving electrical stimulation of the human brain, including those for epilepsy.

## Acknowledgements

We gratefully acknowledge support from the Department of Neurosurgery at the University of Utah.

## Supplementary material

Supplementary material is available at *Brain* online.

### FIGURES

1. Stimulation Guided User Interface

2. K-means Cluster Waveform Classification

3. Cell Type Clusters for Different Data Segregations

4. Distance. Breakdown of Sigmoidal Curve, R2 vs distance for significant cells

5. Examples of Unit Responses to Single Pulse Electrical Stimulations

6. Regression Analysis for Neuronal Features

### TABLES

1. Patient Demographic Information

2. Electrode and Unit Location

3. Clinical Interpretation of Seizure Onset Zone

