## Supplementary Materials for "Cell-type specific responses to single-pulse electrical stimulation of the human brain"

### SUPPLEMENTARY FIGURES

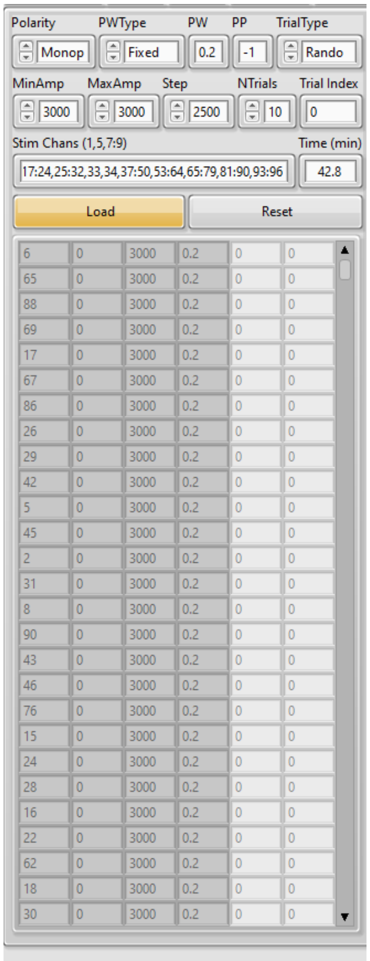


**Supplementary Figure 1. Stimulation Guided User Interface.** Screenshot of the stimulation control guided user interface used to implement the SPES protocol. *Parameters:* Polarity = Monopolar; Pulse-Width Type = Fixed; Pulse-Width = 0.2; Trial Type = Random; = Minimum Amperage = 3000; Maximum Amperage = 3000; Number of Trials = 10; Stim Channels = individualized patient channels; Time = run time for protocol.

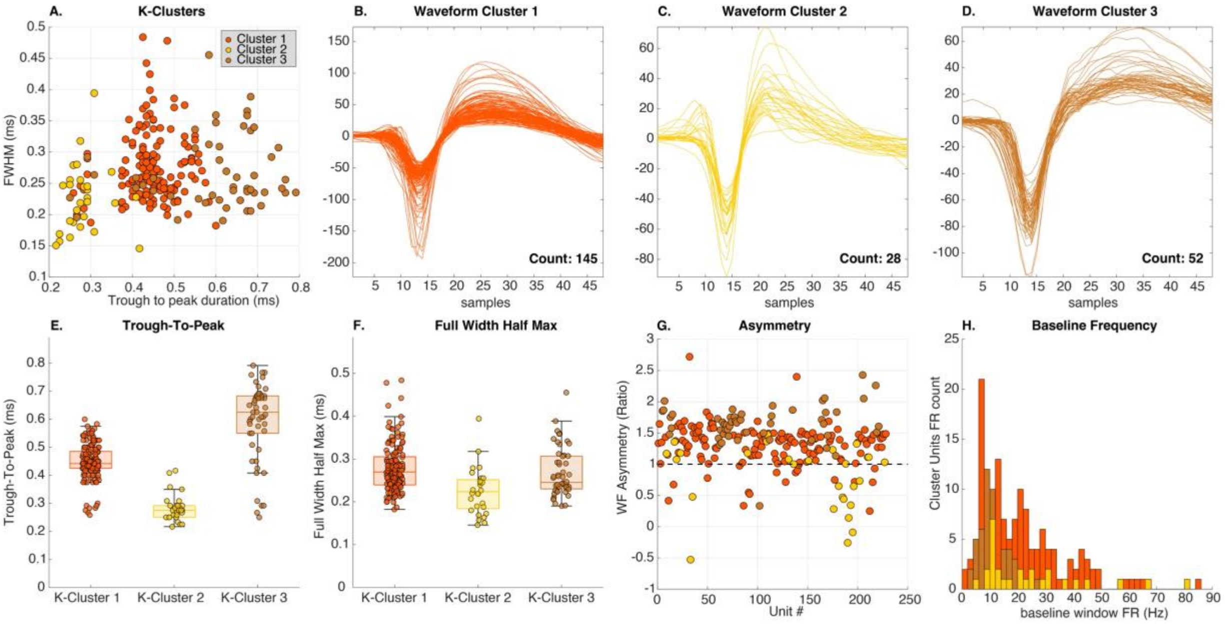


**Supplementary Figure 2. K-means Cluster Waveform Classification. (A)** Cell Types clustered based on K-Means (*k = 3*) calculation **(B)** Cluster K1 waveforms, based on subpanel A orange cluster**. (C)** Cluster K2 waveforms, based on subpanel A yellow cluster**. (D)** Cluster K3 waveforms, based on subpanel A brown cluster**. (E)** Trough-To-Peak (ms) metric for the three K-Clusters (k1 = 0.448 ± 0.064; k2 = 0.283 ± 0.049; k3 = 0.594 ± 0.138) **(F)** Full-Width Half-Max (ms) metric for the three K-Clusters (k1 = 0.278 ± 0.055; k2 = 0.225 ± 0.055; k3 = 0.268 ± 0.057) **(G)** Asymmetry (Ratio) for the three K-Clusters (k1 = 1.305 ± 0.320; k2 = 0.788 ± 0.527; k3 = 1.649 ± 0.309) **( (H)** Baseline Frequency (Hz) (-1.1s -0.1s before stimulation) for the three K-Clusters (k1 = 11.894 Hz ± 6.746; k2 = 24.007 Hz ± 18.771; k3 = 21.511 Hz ± 15.633).

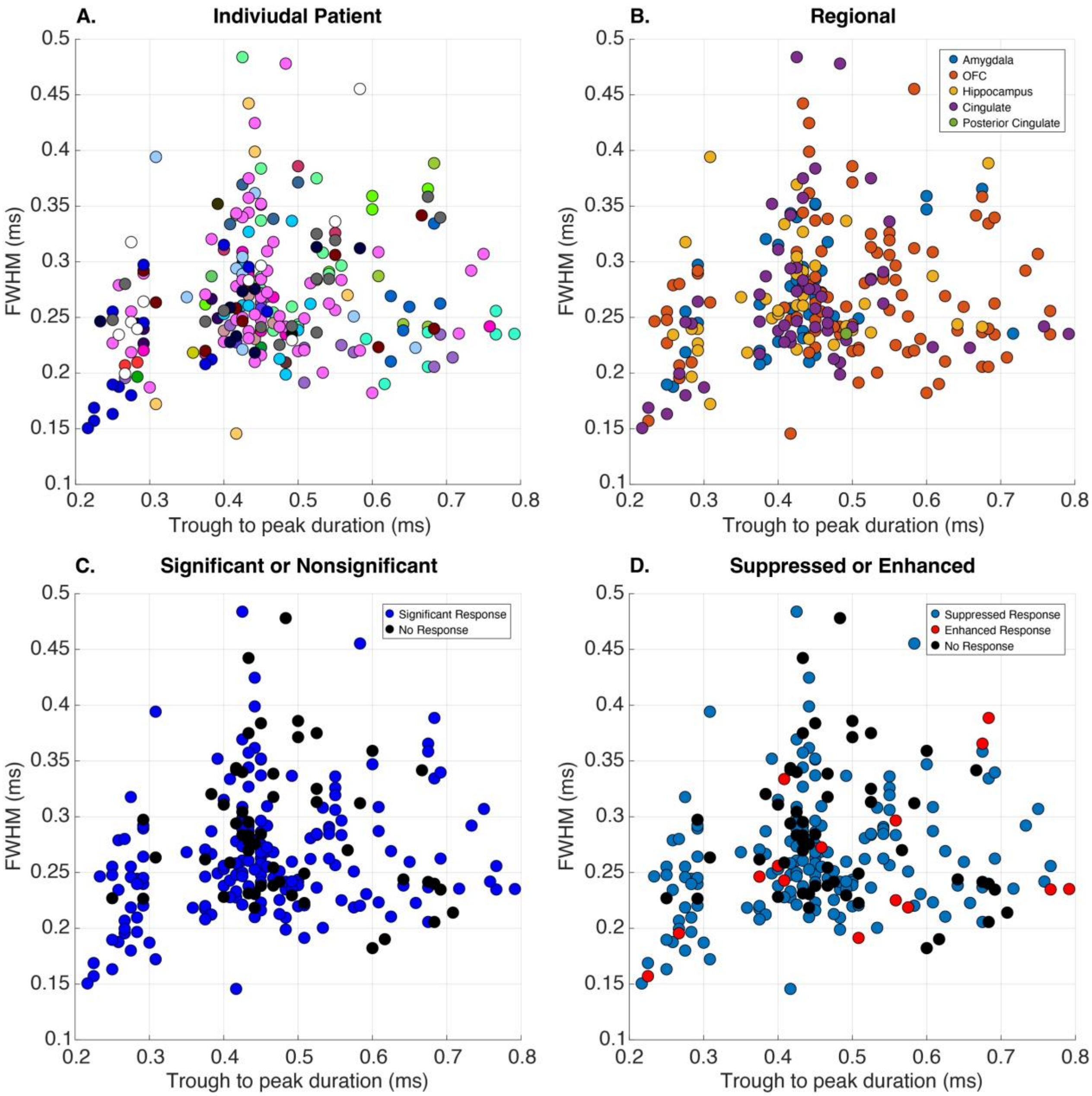


**Supplementary Figure 3. Cell Type Clusters for Different Data Segregations. (A)** Units clustered based on individual patients (individual colors per patient) **(B)** Units clustered based on brain region **(C)** Units clustered based on whether the unit was had significantly changed its firing rate pre-post stimulation (blue = significant, black non-significant) **(D)** Units clustered based on whether the unit showed suppression (blue), enhancement (red), or no change (black) in response

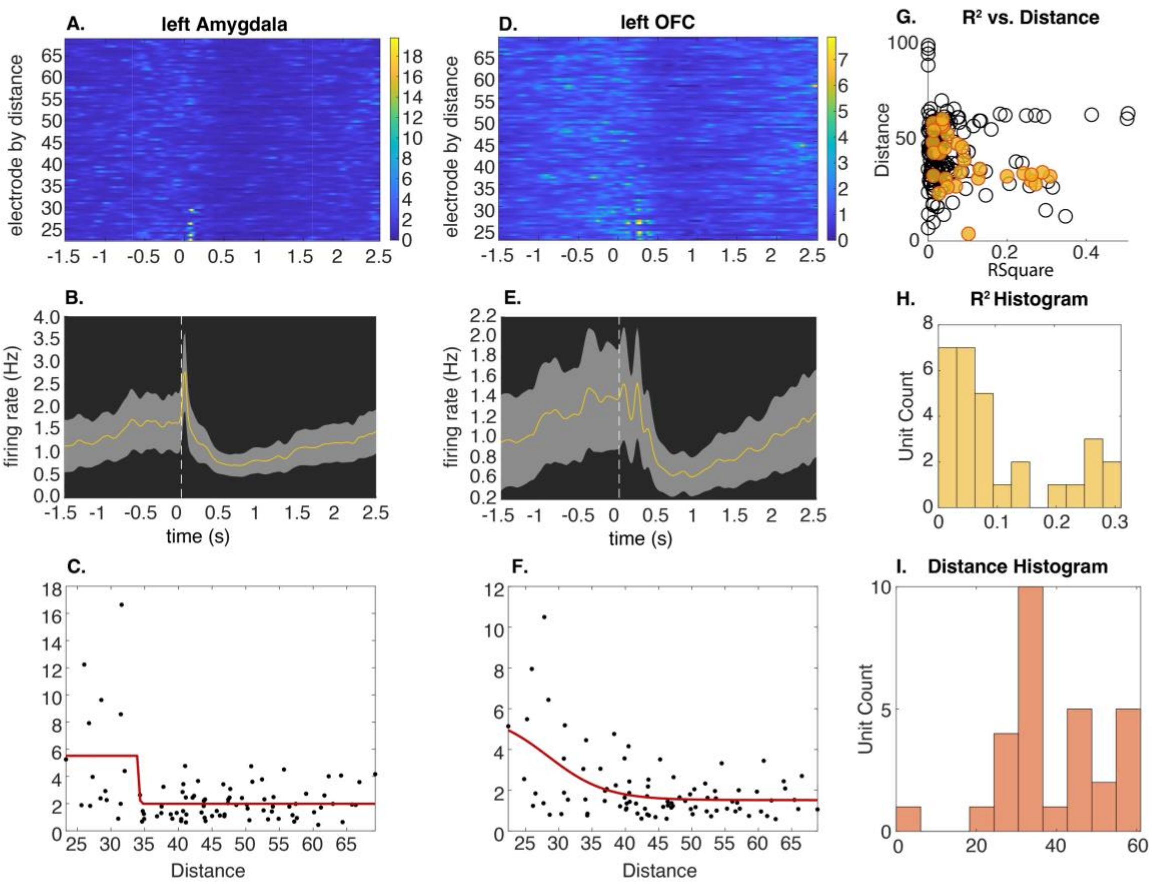


**Supplementary Figure 4. Distance. Breakdown of Sigmoidal Curve, R2 vs distance for significant cells.** A-C, Left Amygdala example. (a) Firing rate of unit ordered by the stimulation electrode by distance over time (stimulation at time zero). (b) Average firing rate over time (stimulation at time zero). (c) Sigmodal curve to determine threshold for evoked firing threshold. D-F, Left OFC example. (d) Firing rate of unit ordered by the stimulation electrode by distance over time (stimulation at time zero). (e) Average firing rate over time (stimulation at time zero). (f) Sigmodal curve to determine threshold for evoked firing threshold. (g) R-squared by distance for significantly evoked firing units (orange) and nonsignificant units (black). (h) Histogram for R-squared values by uit count. (i) Unit count and average distance for evoked firing rates (average 39mm from

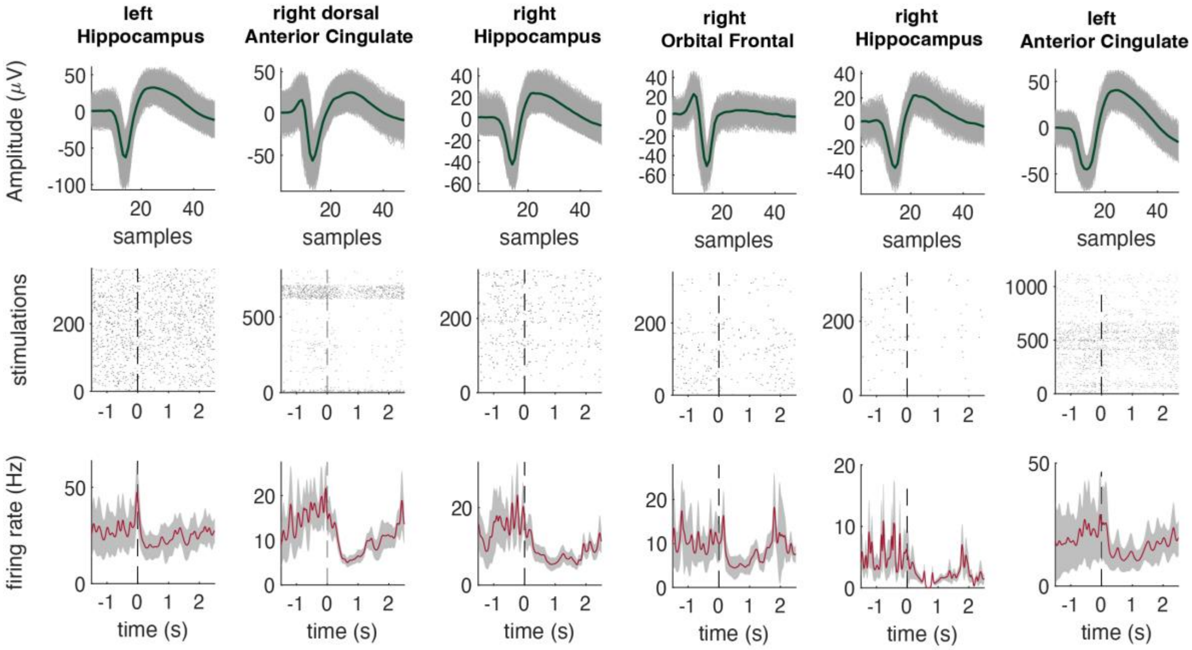


**Supplementary Figure 5. Examples of Unit Responses to Single Pulse Electrical Stimulations.** (*top*) Action potential waveforms. Green line represents mean waveform for each unit. Grey lines represent all waveforms (*middle*) Stimulation raster. Each gray dot represents a stimulation trial (*bottom*) Firing rate average across all trials (mean = red line) with standard deviation (grey). Time zero represents the time of stimulation.

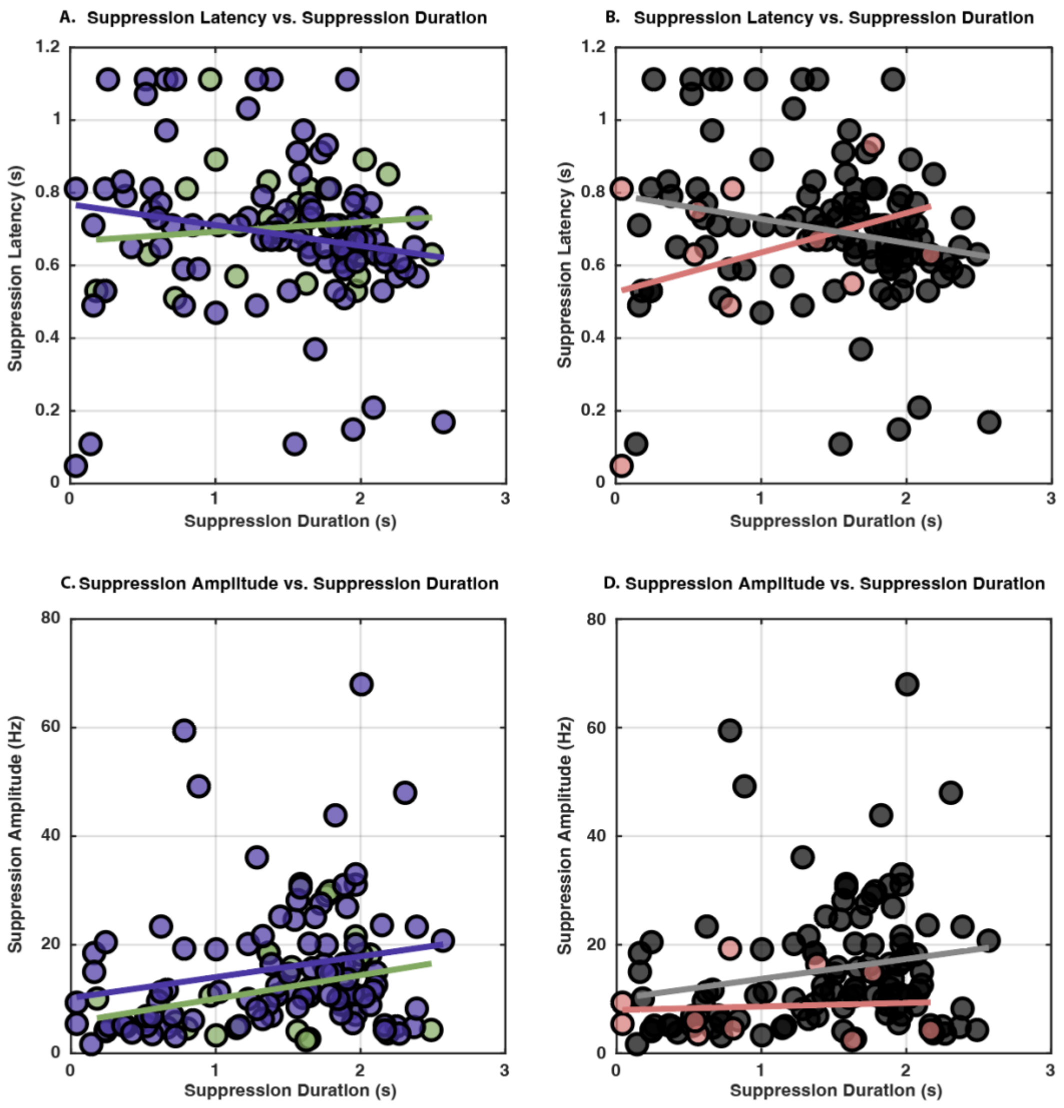


**Supplementary Figure 6. Regression Analysis for Neuronal Features.** (a) Cell type regression between suppression latency (ms) and duration (ms) for interneurons (green) ($t(24) = 0.55$, $p = 0.59$, $R^2 = 0.012$) and principal cells (purple), ($t(90) = -1.661$, $p = 0.100$, $R^2 = 0.03$). (b) SOZ regression between suppression latency (ms) and duration (ms) for within-SOZ (pink) ($t(8) = 0.98$, $p = 0.36$, $R^2 = 0.106$) and outside-SOZ (black) ($t(106) = -2.189$, $p = 0.031$, $R^2 = 0.043$). (c) Cell type regression between suppression amplitude (Hz) and duration (ms) for interneurons (green) ($t(24) = 0.1.529$, $p = 0.139$, $R^2 = 0.089$) and principal cells (purple) ($t(90) = 2.018$, $p = 0.047$, $R^2 = 0.043$). (d) SOZ regression between suppression amplitude (Hz) and duration (ms) for within-SOZ (pink) ($t(8) = 0.225$, $p = 0.828$, $R^2 = 0.006$) and outside-SOZ (black) ($t(106) =2.041$, $p = 0.044$, $R^2 = 0.038$). For all graphs, lines represent respectively regression for each unit group (cell type, SOZ units) with underlaid scatter plots for each unit.

### SUPPLEMENTARY TABLES

**Table 1 Participant Demographic Information**

| Patient # | Gender Identity | Age in Years (At Time of Testing) |
|---|---|---|
| 1 | Male | 26 |
| 2 | Male | 56 |
| 3 | Male | 40 |
| 4 | Female | 42 |
| 5 | Female | 38 |
| 6 | Male | 24 |
| 7 | Male | 35 |
| 8 | Female | 36 |
| 9 | Female | 40 |
| 10 | Male | 24 |
| 11 | Female | 30 |
| 12 | Male | 23 |
| 13 | Male | 43 |
| 14 | Female | 34 |
| 15 | Male | 34 |
| 16 | Female | 40 |
| 17 | Female | 35 |
| 18 | Female | 19 |
| 19 | Female | 23 |
| 20 | Female | 66 |
| 21 | Female | 44 |
| 22 | Female | 42 |
| 23 | Female | 42 |
| 24 | Female | 40 |
| 25 | Female | 34 |
| 26 | Male | 32 |
| 27 | Male | 23 |
| 28 | Male | 49 |
| 29 | Male | 49 |
| 30 | Male | 28 |

Gender Identity and Age in Years at the time of testing (n = 30, 16 females, 36 ± 10.39 years of age

**Table 2 Electrode and Unit Location**

| Patient # | # of Implanted Microelectrodes | Microelectrode Anatomical Location | # of Units | Reference, Ground Electrode Location |
|---|---|---|---|---|
| 1 | 2 | [left Amygdala; left Anterior Cingulate] | [0;1] | Left Orbital Frontal Cortex 6, Bovie pad |
| 2 | 3 | [left medial Orbital Gyrus; left Anterior Cingulate; left Amygdala] | [2;1;4] | Left Insula 9, Bovie pad |
| 3 | 2 | [left Hippocampus; left dorsal Anterior Cingulate] | [1;0] | Right Amygdala 6, Bovie pad |
| 4 | 2 | [left subcallosal area (vmPFC); left Anterior Hippocampus] | [1;0] | Left Insula 8, Bovie pad |
| 5 | 2 | [right Gyrus Rectus; right dorsal Anterior Cingulate] | [0;1] | Left Insula 10, Bovie pad |
| 6 | 2 | [right OFC; right Hippocampus] | [0;4] | Right Insula, Bovie pad |
| 7 | 2 | [right OFC; right Hippocampus] | [2;0] | Right Insula, Bovie Pad |
| 8 | 2 | [left OFC; right Hippocampus] | [0;6] | Left VC 7, Bovie pad |
| 9 | 2 | [left subgenual Cingulate; left Anterior Cingulate] | [2;0] | Left Insula 10, Bovie pad |
| 10 | 2 | [left OFC; left subgenual Cingulate] | [2;3] | Right Hippocampus 9, Bovie pad |
| 11 | 2 | [right OFC; right Hippocampus] | [4;1] | Left CV 8, Bovie pad |
| 12 | 2 | [right OFC; right Hippocampus] | [2;2] | Right RCD 7, Bovie Pad |
| 13 | 4 | [left OFC; left ventral Cingulate; left dorsal Anterior Cingulate; right Anterior Hippocampus] | [8;8;0;0] | Left CM 9, Bovie pad |
| 14 | 3 | [right OFC; right dorsal Anterior Cingulate; left Anterior Hippocampus] | [0;2;2] | Right RCH 7, Bovie pad |
| 15 | 2 | [right OFC; right Hippocampus] | [6;0] | Right VCG 14,13 |
| 16 | 3 | [right ventral Cingulate; left ventral Cingulate, left OFC] | [0;1;5] | Right DCG 4, Bovie pad |
| 17 | 2 | [right OFC; right ventral Cingulate] | [9;2] | Right OFC 6, Bovie pad |
| 18 | 3 | [right OFC; right dorsal Anterior Cingulate; right Anterior Hippocampus] | [5;1;1] | Right Insula 9,10 |
| 19 | 3 | [right Anterior Cingulate; right Anterior Hippocampus; left Hippocampus] | [3;2;5] | Left Insula 8,9 |
| 20 | 3 | [right Hippocampus; left Anterior Hippocampus; left Anterior Cingulate] | [0;0;2] | Right CMT 7,8 |
| 21 | 3 | [left OFC; left mid Cingulate; right Hippocampus] | [2;2;5] | Left Insula 8,9 |
| 22 | 3 | [left OFC; left Anterior Cingulate; right Amygdala] | [10;2;9] | Left Anterior Insula 8,9 |
| 23 | 3 | [left OFC; left Anterior Cingulate; right Amygdala] | [12;4;15] | Left Anterior Insula 8,9 |
| 24 | 3 | [left OFC; left ventral Cingulate; right Amygdala] | [5;2;3] | Left Anterior Insula 7,8 |
| 25 | 1 | [right Posterior Cingulate] | [1] | Right posterior Insula 7,8 |
| 26 | 3 | [right OFC; right Anterior Cingulate; left Hippocampus] | [0;0;1] | Right RCM 9,10 |
| 27 | 3 | [left OFC; left Anterior Cingulate; right Hippocampus] | [5;5;0] | Left Anterior Thalamus 9,10 |
| 28 | 3 | [left OFC; left Anterior Cingulate; left Amygdala] | [4;6;8] | Right posterior Insula 7,8 |
| 29 | 3 | [left OFC; right Hippocampus; left Anterior Cingulate] | [8;4;2] | Left Insula 7,8 |
| 30 | 3 | [right OFC; right Anterior Cingulate; right Anterior Hippocampus] | [1;6;4] | Right Anterior Insula, 9,10 |

Descriptive for each patient from columns left to right. The corresponding patient ID, number of implanted microelectrodes, location of microelectrodes electrodes based on Neuromorphometrics Atlas from LeGUI, number of units in sorted in each of those locations, and reference and ground electrode locations.

**Table 3 Clinical Interpretation of Seizure Onset Zone**

| Patient # | Behnke-Fried Electrode Locations | Clinical Seizure Onset Zone |
|---|---|---|
| 1 | [left Amygdala; left Anterior Cingulate] | left FOP |

| | | |
|---|---|---|
| 2 | [left medial Orbital Gyrus; left Anterior Cingulate; **left Amygdala**] | left anterior hippocampus; left amygdala |
| 3 | [**left Hippocampus**; left dorsal Anterior Cingulate] | hippocampal (left dominance) |
| 4 | [left subcallosal area (vmPFC); **left Anterior Hippocampus**] | left anterior hippocampus; spread to left posterior hippocampus and amygdala. (left hippocampus main) |
| 5 | [right Gyrus Rectus; right dorsal Anterior Cingulate] | left hippocampus; right hippocampus |
| 6 | [right OFC; **right Hippocampus**] | right hippocampus; right amygdala |
| 7 | [right OFC; **right Hippocampus**] | right hippocampus; right amygdala; left hippocampus; left amygdala |
| 8 | [left OFC; right Hippocampus] | left hippocampus |
| 9 | [left subgenual Cingulate; left Anterior Cingulate] | right hippocampus; right amygdala |
| 10 | [left OFC; left subgenual Cingulate] | left hippocampus |
| 11 | [right OFC; **right Hippocampus**] | right hippocampus; orbital frontal gyrus and insula |
| 12 | [right OFC; right Hippocampus] | right PO and right CM |
| 13 | [left OFC; left ventral Cingulate; left dorsal Anterior Cingulate; right Anterior Hippocampus] | left insula; left hippocampus |
| 14 | [**right OFC**; right dorsal Anterior Cingulate; left Anterior Hippocampus] | right amygdala; right anterior hippocampus; right orbital frontal gyrus (subclinical) |
| 15 | [right OFC; **right Hippocampus**] | right anterior and posterior hippocampus; right orbital frontal gyrus (spread) |
| 16 | [right ventral Cingulate; left ventral Cingulate, left OFC] | right hippocampus; right amygdala; left anterior hippocampus |
| 17 | [right OFC; right ventral Cingulate] | right anterior hippocampus; right amygdala |
| 18 | [right OFC; right dorsal Anterior Cingulate; **right Anterior Hippocampus**] | right amygdala; right anterior/posterior hippocampus |
| 19 | [right Anterior Cingulate; right Anterior Hippocampus; left Hippocampus] | left amygdala |
| 20 | [right Hippocampus; left Anterior Hippocampus; left Anterior Cingulate] | bilateral hippocampi and amygdala |
| 21 | [left OFC; left mid Cingulate; right Hippocampus] | left hippocampus |
| 22 | [left OFC; left Anterior Cingulate; right Amygdala] | left anterior hippocampus, left posterior hippocampus |
| 23 | [left OFC; left Anterior Cingulate; right Amygdala] | left anterior hippocampus, left posterior hippocampus |
| 24 | [left OFC; left ventral Cingulate; right Amygdala] | left hippocampus; left amygdala |
| 25 | [right Posterior Cingulate] | heschel's gyrus |
| 26 | [right OFC; right Anterior Cingulate; left Hippocampus] | right hippocampus and amygdala |
| 27 | [left OFC; left Anterior Cingulate; right Hippocampus] | left hippocampus; left amygdala |
| 28 | [left OFC; left Anterior Cingulate; left Amygdala] | right anterior hippocampus; left hippocampus |
| 29 | [left OFC; **right Hippocampus**; left Anterior Cingulate] | left hippocampus; left amygdala; right hippocampus; right amygdala |
| 30 | [right OFC; **right Anterior Cingulate**; right Anterior Hippocampus] | right anterior cingulate |

Details of patient ID, Behnke-Fried Electrode Locations (microelectrodes placed within SOZ bolded), and the clinically defined seizure onset zone by attending Epileptologist.
